# Regulation of brassinosteroid signaling through Arabidopsis transcription factor TCP8 by a conserved post-translational mechanism

**DOI:** 10.1101/2025.01.30.635591

**Authors:** Benjamin J. Spears, Samuel A. Magnabosco, Raegan Mozal, Nyle M. Beauford, Noah C. Baker, Quinn Voeglie, Walter Gassmann

## Abstract

The ability of plants to respond to changing environmental conditions is dependent on their capacities to produce fast, transient, and reversible changes to stress-responsive gene expression profiles. Post-translation modification of transcription factors has been identified as a contributing mechanism to these responses but has yet to be explored mechanistically in the TCPs, a plant-specific TF family known to be broadly involved in phytohormone signaling governing biotic and abiotic stress responses. Here, we use a recombinant expression system and proteomic approach to identify the specific residues at which AtTCP8 is covalently modified by the small peptide SUMO and demonstrate that loss of key lysine residues abolishes modification of the TF. We validate these observations by demonstrating the failure of a sumoylation-site-deficient AtTCP8 mutant to complement a known *tcp8* BR signaling deficiency phenotype natively in *Arabidopsis* and reduced transcriptional activity of the protein at a known regulatory target promoter. Further, we show that class I TCP orthologs in the model non-vascular plant *Physcomitrium patens* are sumoylated as well in our system, a novel finding for TFs in this organism. These data support a conserved model for post-translational regulation by SUMO of phytohormone-responsive gene expression between vascular and non-vascular plants.

## Introduction

Plants produce, perceive, and respond to a variety of chemical signals to allow the optimization of their growth and development under increasingly variable environmental conditions. A major phytohormone signaling pathway controlling these processes in plants involve a group of plant-specific steroids known as the brassinosteroids (BRs). The pathway itself is well-defined in *Arabidopsis*, and begins with the extracellular perception of the hormone by plasma membrane-localized receptors such as AtBRI1 (Li and Chory, 1997; Li *et al*., 2002). Internalization of signal results in the ubiquitination and subsequent deactivation of the regulatory GSK3-like kinase AtBIN2 which under BR-deplete conditions suppressively phosphorylates key transcription factors (TFs) BZR1 and BZR2/BES1 (He *et al*., 2002; Zhao *et al*., 2002). The newly dephosphorylated TFs are then able to enter the nucleus and regulate the expression of genes involved in growth and development (Yin *et al*., 2002; Ryu *et al*., 2007). Their activity is further balanced through cooperation with a variety of other non-canonical bHLH TFs including members of the PIF and TCP families, a feature of this regulatory system that may allow for complex arrangements of transcriptional output to fine-tune a plant’s response to stimuli (Oh *et al*., 2012; Spears *et al*., 2022; Diao *et al*., 2023) These combinatorial TF interactions are not as well defined and represent a broader point of control for regulatory interactions across phytohormone signal transduction pathways (Zhang *et al*., 2013; Pfeiffer *et al*., 2014). Post-translational modification of molecular components at multiple steps is a key element of BR and most other phytohormone signaling pathways (Withers and Dong, 2017; Cui *et al*., 2024; Mao *et al*., 2024), and is a likely mechanism involved in establishing TF dimerization patterns involved.

The TCP family has been characterized as signaling pathway components of multiple phytohormone signaling pathways in both vascular and nonvascular plants (Wang *et al*., 2022), with diverse functions identified within broad categories of growth, development, and biotic and abiotic stress responses (Guo *et al*., 2010; Danisman *et al*., 2012; Wang *et al*., 2015; Gonzalez-Grandio *et al*., 2017) . The approximately 60 amino acid TCP domain is hallmark feature of the 24-member family in *Arabidopsis* and is critical to DNA-binding and dimerization capacities of each protein (Nicolas and Cubas, 2016). TCPs are sub-categorized into Class I or Class II on the basis of small variations within and around that N-terminal domain. Interestingly, TCPs often share very little sequence identity with each other outside of the TCP domain, which aligns with high levels of intrinsic disorder and additional prion-like domains commonly observed in the family that resemble those found in other highly disordered and phytohormone-responsive TFs like Auxin Response Factors (ARFs). Intrinsically disordered regions (IDRs) of TFs tend to be sites of post-translational modification which offer temporary stabilization of structure in an environmentally contextual and reversible manner, and thus the achievement of heterodimerization patterns that were not previously favored (Hill, 2015; Salladini *et al*., 2020). In this way, the modification of TCPs may be key to their diverse functions in plant physiology, or the ability of a single protein to be able to achieve antagonistic signaling behavior across pathways.

Class I *Arabidopsis* TCP8 was previously shown to be sumoylated (Mazur *et al*., 2017), a modification involving the small SUMO peptide and activities of an E1-E3 enzyme cascade at lysine residues and that commonly alters subcellular localization, dimerization patterns, and functional activities of target substrates. Closely-related TFs AtTCP14 and AtTCP15 were also shown to be sumoylated, and have been further characterized as targets of the Arabidopsis OGT protein AtSPINDLY to modulate their roles in cytokinin signaling (Steiner *et al*., 2012, 2016), and the MAP kinase AtMKP8 in gibberellin-related dormancy (Zhang *et al*., 2019a) .

AtTCP8 has been identified as a transcriptional regulator of AtBZR1 and AtBZR2, and a heterodimerization partner of AtBZR2 in its role in promoting BR signaling (Spears *et al*., 2022); interestingly, AtBZR1 and AtBZR2 have also been identified as sumoylated proteins, with modification promoting or repressing their activities BR signaling activities, respectively (Zhang *et al*., 2019b; Srivastava *et al*., 2020). We hypothesized that like AtBZR2, the sumoylation of AtTCP8 is significant to its function as a regulator of phytohormone signaling networks. In this study, we identify the specific residues at which AtTCP8 is post-translationally modified by SUMO and using protein-protein interaction and promoter transactivation assays in tobacco characterize altered behaviors of a sumoylation-null mutant of AtTCP8. We provide data that demonstrate the sumoylation of AtTCP8 to be essential to its known function as positive regulator of BR signaling. We further establish *Physcomitrium patens* class I TCPs as one of the first described sumoylated proteins in bryophytes, suggesting that this regulatory mechanism is ancestrally conserved between vascular and nonvascular plants as an important molecular switch to govern complex TF activities controlling plant growth, development, and stress responses.

## Methods

### Plant materials

Arabidopsis plants were grown on ½ strength MS 0.8% phytoagar plates under an 8-hr light and 16-hour dark cycle at 24 °C, 70-80 % RH, and a light intensity of 140-180 µM photons sq m^-1^/ sec^-1^. *Nicotiana benthamiana* plants used in transactivation and interaction assays were grown under an 8-hour light and 16-hour dark cycle at 22 °C.

The transgenic *pAtTCP8:TCP8-HA* (TCP8-HA) and *tcp8 tcp14 tcp15* (*t8t14t15*) triple mutant lines have been previously characterized (Kim *et al*., 2014; Spears *et al*., 2019, 2022) and transgenic sumoylation-site mutant *pAtTCP8:TCP8^K57,230R^-HA* lines were generated in the *t8t14t15* background by floral dipping (Clough and Bent, 1998). Two independent representative lines were selected and evaluated in this study.

### Molecular Cloning

*pBZR2* promoter constructs in the vector pYXT1, *35s:HA-TCP8* expression constructs in the vector pBA-HA, assorted split-luciferase constructs, and Arabidopsis complementation construct *pTCP8:TCP8-HA* in the pMDC83 vector have previously been published (Kim *et al*., 2014; Spears *et al*., 2019, 2022).

Full-length cDNAs of *PpTCP1*, *PpTCP2*, and *PpTCP4* were amplified from cDNA template synthesized from *P. patens* cv. *Grandsen* gDNA and cloned into the Gateway-compatible donor vector pDONR221 (Invitrogen). Donor clones were used as a template for PCR-based addition of HindIII and EcoRI restriction sites to the CDS. Products were digested and ligated into corresponding restriction sites in the His/T7-tag vector pET-28a. The full-length cDNA of *AtTCP8* was similarly cloned into the GST-tag vector pGEX-4T3.

All single and higher-order *KxR* mutant constructs in this study (*AtTCPs* and *PpTCPs*) were generated by repeated site-directed mutagenesis of the existing His-T7, HA, and nLUC/cLUC-tagged constructs, according to standard protocol. Primers used have been included in supplemental table S1.

### LC-MS/MS identification of AtTCP8 sumoylation sites

#### Sequential Purification

Fresh transformants of *E. coli strain BL21 (DE3)-pLysS* expressing GST-AtTCP8, HIS-T7 SUMO1-RGG (Miller *et al*., 2010), and sumoylation machinery (Okada *et al*., 2009) were grown in 400 ml culture under selection at 25 °C overnight. Pelleted culture was resuspended in 25 ml cold TBS with bacterial protease inhibitor and lysed by French press before centrifugation at 14,000 x *g*. Lysate was divided into two fractions, each diluted to 13 mL with cold TBS. Glutathione agarose beads were prepared according to manufacturer’s instructions (G-Biosciences, St. Louis MO USA). 1 mL of beads were added to each fraction and incubated shaking at 4 °C overnight. Beads were washed with cold TBS before eluting captured protein with 500 µL 10 mM glutathione for 15 min. For each replicate, the first 3 eluted fractions were combined and diluted with 1/3 volume of denaturing Ni-NA lysis buffer. 200 µL prepared Ni-NTA bead slurry was added to 7 mL of the diluted lysate and incubated on a rotating shaker at 4 °C overnight. Beads were batch-watched 3 times before resuspension in Ni-NTA lysis buffer including 6 M urea. Identical preparations were made for GST-AtTCP15.

#### Preparation

Protein samples were thawed and reduced with DTT, alkylated with IAA, and digested with 4 µg trypsin at 37 °C overnight. Samples were acidified after double digest, lyophilized to approximately 2 mL, and subjected to C18 tip peptide enrichment and purification. Following elution from the C18 tips, an equal amount of Milli-Q water was added to dilute samples to 35 % ACN, 0.5 % formic acid and lyophilized.

Samples were resuspended in 5 % acetonitrile/ 1 % formic acid and transferred to an autosampler vial. Samples were loaded into a C8 trap column (ThermoFisher, µ-precolumn-300 µm i.d. x 5mm, C8 Pepmap 100, 5 µm, 100 Å). Peptides were eluted onto a 25 cm, 150 µm i.d. pulled-needle analytical column packed with HxSIL C18 reversed phase resin (Hamilton Co.) Peptides were separated and eluted with a continuous gradient of acetonitrile at 400 nL/min for analysis on an attached LTQ Orbitrap XL mass spectrometer.

#### LC-MS/MS Conditions

Initial conditions were 5% of solution B (99.9% acetonitrile, 0.1% formic acid), followed by 2 min at 5% B. Gradient to 20 % B for 45 min. Gradient from 20 % B to 30 % B in 50 min. Gradient 30- 90 % B over 15 min. Hold at 90 % B for 22 min. Ramp back to and hold at initial conditions for 5 min.

FTMS data were collected (30,000 resolution, 1 microscan, 300-1800 m/z, profile) and then for each 3 second cycle the 9 most-abundant peptides (ignore +1 ions, >2,000 counts) were selected for MS/MS (2 m/z mass window, 35% normalized collision energy, centroid). Spray voltage applied was 1.6 kV.

Sorcerer-Sequest and MASCOT were used to conduct database searches of raw data in Swissprot/Uniprot, with SUMO (R-GG after trypsin digestion) allowed as a variable modification in both searches.

#### Phylogenetic analysis

Putative and characterized SUMO proteins and E1/E2/E3 enzymes for *P. patens*, *A. thaliana*, *Z. mays*, *G. Max*, and *S. lycopersicum* were datamined and phylogenetic trees produced using Geneious. Briefly, alignments were performed under default setting options for global alignment with free end gaps and the Biosum62 cost matrix. Interactive Tree of Life was used to enhance presentation of the Geneious output model.

#### Arabidopsis BL sensitivity assay

Stratified seed was plated on square ½ MS plates treated with either DMSO or 100 nM 24-epiBL (Sigma E1641) in the media. Plates were placed vertically at 4 °C for two days with foil covering before removal of foil and transfer to a short-day growth chamber. Root length was measured using ImageJ after 10 days.

#### Transactivation assay

Assays were performed as previously described (Spears *et al*., 2022). Briefly, the 2 kb BZR2 promoter:GUS construct were transformed with *35S:HA-TCP8* and *35S:HA-TCP8^K57,230R^* and co- inoculated at OD_600_ 0.2 into *N. benthamiana* leaves by syringe. Tissue was collected 48 hours after inoculation with a 1 cm diameter hole punch, 4 leaf discs per sample. Protein was isolated in extraction buffer and GUS quantified according to a standard 4-MUG assay. Protein concentration was determined by Bradford assay and GUS levels calculated before normalization to the empty vector control.

#### Split luciferase protein interaction assay

Assays were performed as previously described (Spears *et al*., 2022). Briefly, nLUC tested constructs (GUS and BZR2) were transformed with AtTCP8-HA-cLUC and AtTCP8^K57,230R^-HA- cLUC into *A. tumefaciens* C58C1 and co-inoculated at OD_600_ 0.2 into *N. benthamiana* leaves by syringe infiltration. 0.5 cm diameter leaf discs were taken after 48 hours and luciferase activity determined with a BioTek Synergy HTX plate reader.

#### *E. coli* sumoylation assay

Assays were performed as previously described (Mazur *et al*., 2017) with modifications. Briefly, AtTCP8, AtTCP8^K57,230R^, and PpTCP1-4 were cloned into pET28a and co-expressed in BL21 (DE3) strain *E. coli* with pACTCDuet:SAE1/SAE2 and pCDFDuet:AtSUMO1/SCE1 as a triple plasmid transformant under antibiotic selection. Colonies were picked and grown to OD_600_ ∼1.0 before inducing protein expression with 0.2 mM IPTG. Cultures were shaken overnight at 20 °C. 500 µL of each culture was pelleted and then resuspended in 100 µL 1x Laemmli buffer and denatured at 95 °C for 5 minutes before loading onto an 8% SDS-PAGE gel. Proteins were separated then blotted onto PVDF membrane (Millipore-Sigma). T7-directed antibody was applied after blocking overnight at 4 °C at a dilution of (1:20,00 antibody:5% milk) and proteins were visualized with chemiluminescence on an LI-COR Odyssey M imager.

#### In silico analyses

Putative sumoylation sites were identified in AtTCP8 and PpTCP1-4 using the GPS-SUMO tool under default regression settings and high stringency threshold (Gou *et al*., 2024). Regions of intrinsic disorder and PLDs were identified using the PLAAC and PrDOS platforms under default settings (Ishida and Kinoshita, 2007; Lancaster *et al*., 2014).

#### Accession and Identifier numbers

The accession and identifier numbers for *A. thaliana* and *P. patens* genes described in this manuscript are as follows: At1g19350 (*BZR2*), At1g58100 (*TCP8*), At1g69690 (*TCP15*), At1g75080 (*BZR1*), At3g47620 (*TCP14*), At4g39400 (*BRI1*), At4g18710 (*BIN2*), At1g18150 (*MPK8*), At3g11540 (SPINDLY), Pp3c10_20400V3.8 (*PpTCP1*), Pp3c3_24660V3.1 (*PpTCP2*), Pp3c3_24450V3.3 (*PpTCP3*), and Pp3c10_14310V3.1 (*PpTCP4*).

## Acknowledgements

Many thanks to JBS and LJC for technical assistance and feedback during the early days of this study, and to CAC for proofreading and sound advice. We are grateful to the Gehrke Proteomics Center at the University of Missouri for their support of the work outlined in this manuscript, the Butler University Holcomb Awards Committee for supporting the undergraduate-driven work in this study, and to ASPB for the continued engagement and support of students at PUIs through diverse funding mechanisms.

## Results

### Identification and validation of AtTCP8 sumoylation sites

In order to identify the specific residues at which AtTCP8 is modified by SUMO, we recombinantly expressed GST-TCP8 in *E. coli* with using a reconstituted sumoylation system (Okada *et al*., 2009) and then sequentially isolated the population of sumoylated TCP8 by Ni-NTA purification. Conjugation of SUMO protein to the target is visualized by higher-molecular weight band shifts in the presence of functional SUMO1 (RGG), but absent in the presence of a SUMO1 isoform lacking the diglycine motif (AA). Purified protein samples were analyzed by LC-MS/MS and sumoylated TCP8 fragments identified with MASCOT, with lysine residues K57, K216, and K230 exhibiting enrichment for the modification (Fig. 1).

**Figure 1.**
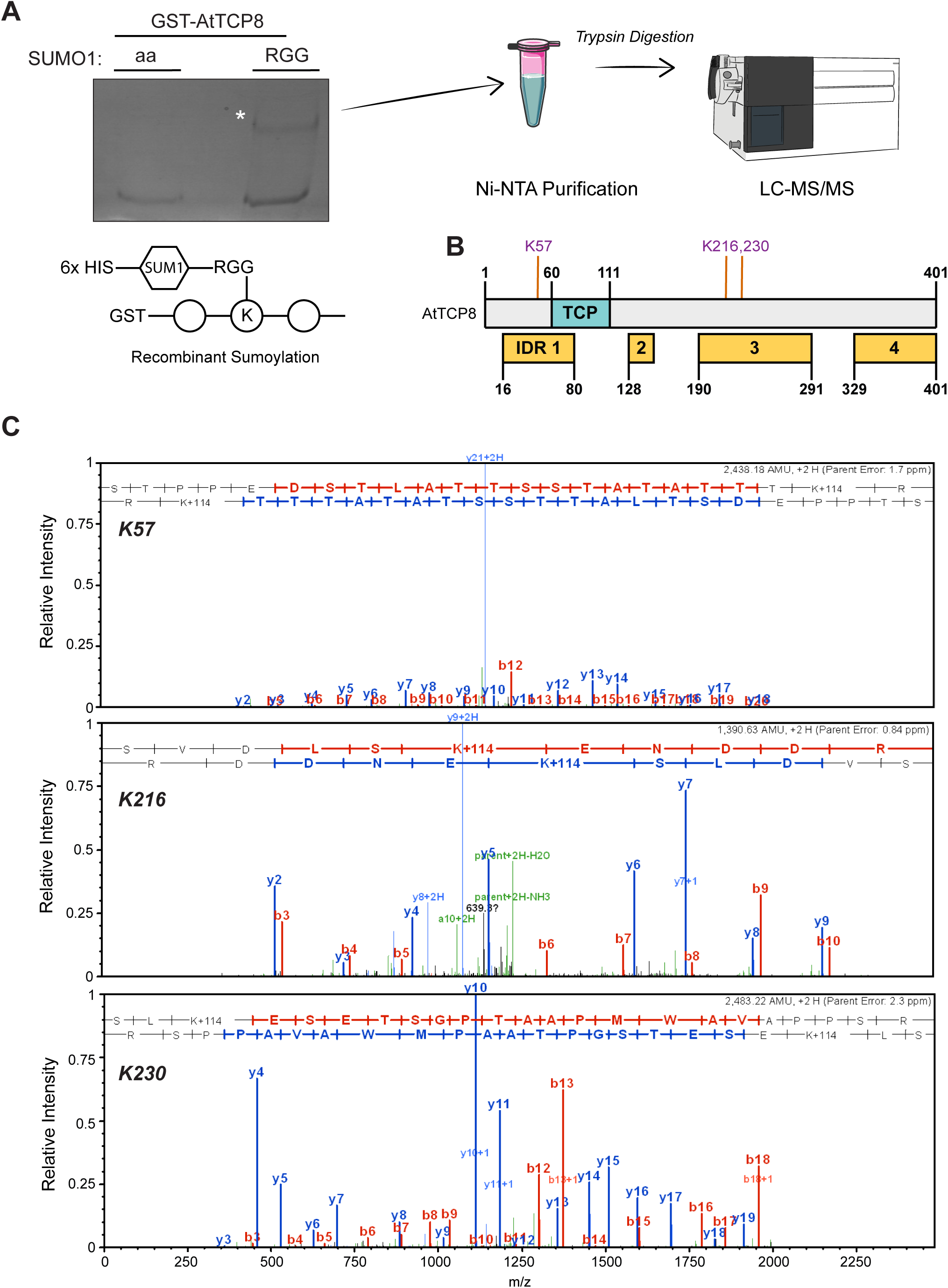
Identification of AtTCP8 sumoylation sites by LC-MS/MS. **A)** AtTCP8 cloned into a GST-tag vector was recombinantly expressed with required sumoylation machinery and modified HIS-tagged AtSUMO1 (RGG) in *E. coli.* SUMO-conjugated moities were purified by Ni-NTA column and digested on-bead with trypsin before LC-MS/MS run. **B)** Protein model of AtTCP8 showing significantly enriched sumoylation events identified at lysines K57, K216, and K230 within disordered regions of the protein. Idenfitied residues are located within regions of predicted intrinsic disorder. **C)** MASCOT was used to identify peptide fragments shown here with observed *y* and *b* ions inrepresentative MS/MS spectra.

As verification of the MS/MS results, we introduced lysine-to-arginine null mutations (KxR) of the GST-TCP8 construct by site-directed mutagenesis to eliminate each identified sumoylation site and then evaluated sumoylation status of each resulting recombinant protein. We focused initial efforts on higher order sumoylation site mutations of both K57 and K230, due to their stronger *in silico* predictions for modification. Although no single KxR mutation exhibited detectable differences in sumoylation comparable to WT AtTCP8 (Fig S1), sumoylation was abolished in the AtTCP8^K57,230R^ double mutant protein as observed by loss of band shift in western blotting (Fig. 2), confirming that lysines K57 and K230 are required for sumoylation of AtTCP8 in the recombinant system. The lack of any additive effect by loss of K216 in the AtTCP8^K57,216,230R^ triple mutant suggests that K57 and K230 may also be sufficient for detectable sumoylation in this system.

**Figure 2.**
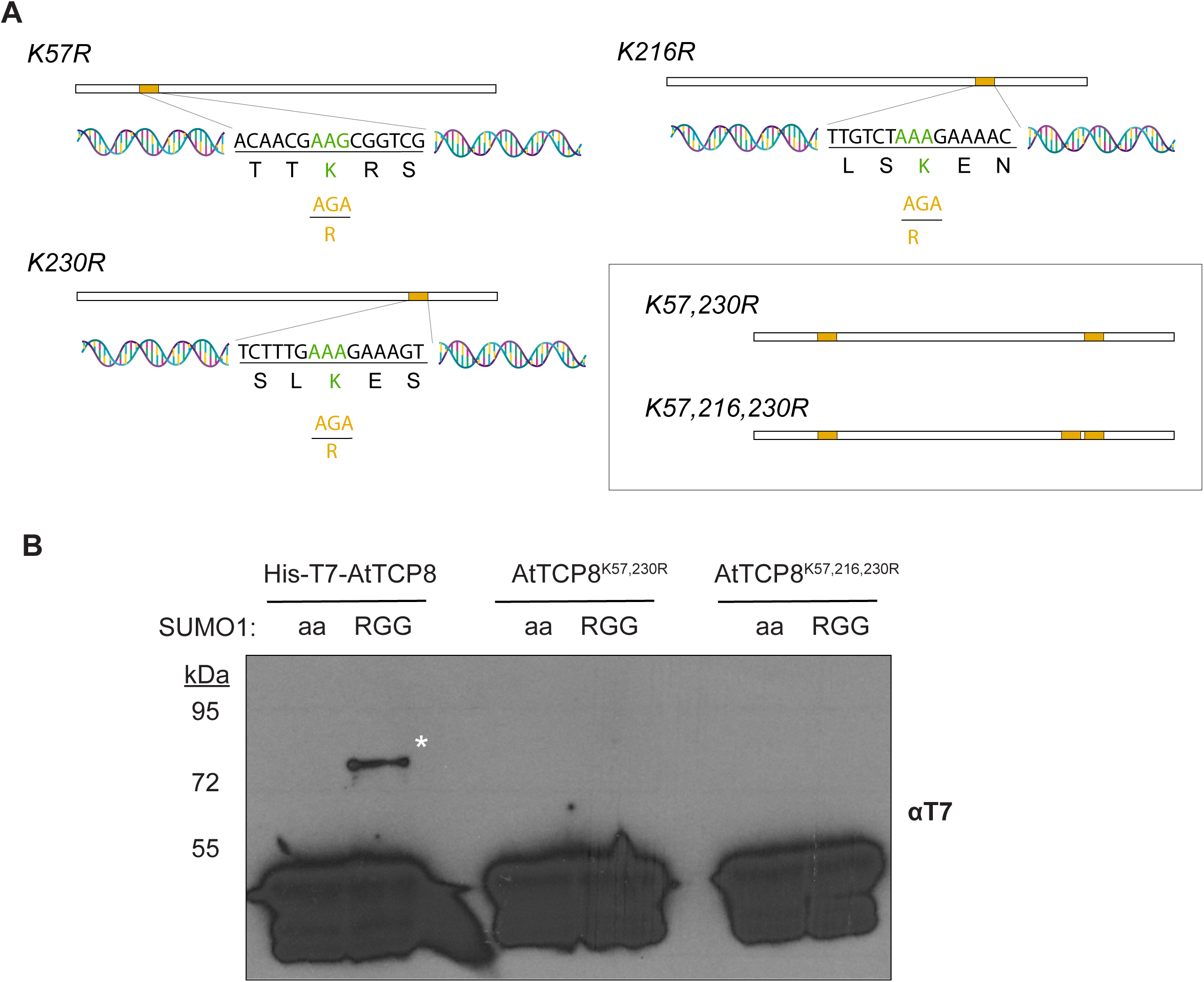
Sumoylation of AtTCP8 *in vitro* is attenuated by K57R and K230R mutations**. A)** Gene models depicting the site-directed mutagenesis of sumoylation-site lysine residues to argenine, including the double and triple mutants AtTCP8^K57,230R^ and AtTCP8^K57,216,230R^. **B)** The K57,230R mutation is sufficient to attenuate the band shift (asterisk) caused by sumoylation of AtTCP8 by HIS-SUMO1-RGG in the recombinant sumoylation system, as seen here with a T7 tag-directed antibody. Similar results were observed in at least three repeat experiments.

### Physcomitrium patens class I TCPs are sumoylated

To date, six TCP-family TFs have been identified in *Physcomitrium patens*, a model non-vascular plant (Ortiz-Ramírez *et al*., 2016). Of these, four are class I, similar to AtTCP8 (Fig. 3A). Like AtTCP8, PpTCP1-4 all exhibit similar patterns of intrinsic disorder, prion-like domains, and multiple predicted sumoylation sites near the N-terminal TCP domain and C-terminal disordered regions (Fig 3B). Although the sumoylation system of *P. patens* is almost completely uncharacterized, enzyme cascade components has been identified sharing sequence similarity with vascular species (Ghosh *et al*., 2024) (Fig. S2). We therefore suspected that sumoylation of AtTCP8 may represent an ancestrally conserved regulatory mechanism identifiable in bryophytic PpTCPs. To explore this possibility, we cloned coding sequences of three class I *PpTCPs* from *P. patens* cv. *Grandsen* and tested whether they could be sumoylated in the recombinant system; however, we were unable to successfully clone *PpTCP3*. We identified sumoylated band shifts for PpTCP1, PpTCP2, and PpTCP4 at a similar increase in molecular weight as for AtTCP8 (Fig. 4A). To verify this finding for a representative member, we used site-directed mutagenesis to generate a *PpTCP1^K279R^* mutant construct for its most highly predicted sumoylation site. Sumoylation of the PpTCP1^K279R^ mutant was strongly reduced, suggesting both that the residue in question is necessary for sumoylation of PpTCP1 in our system (Fig 4B) and that the post-translational modification of class I TCPs near their disordered regions by SUMO may be conserved between vascular and non-vascular plants.

**Figure 3.**
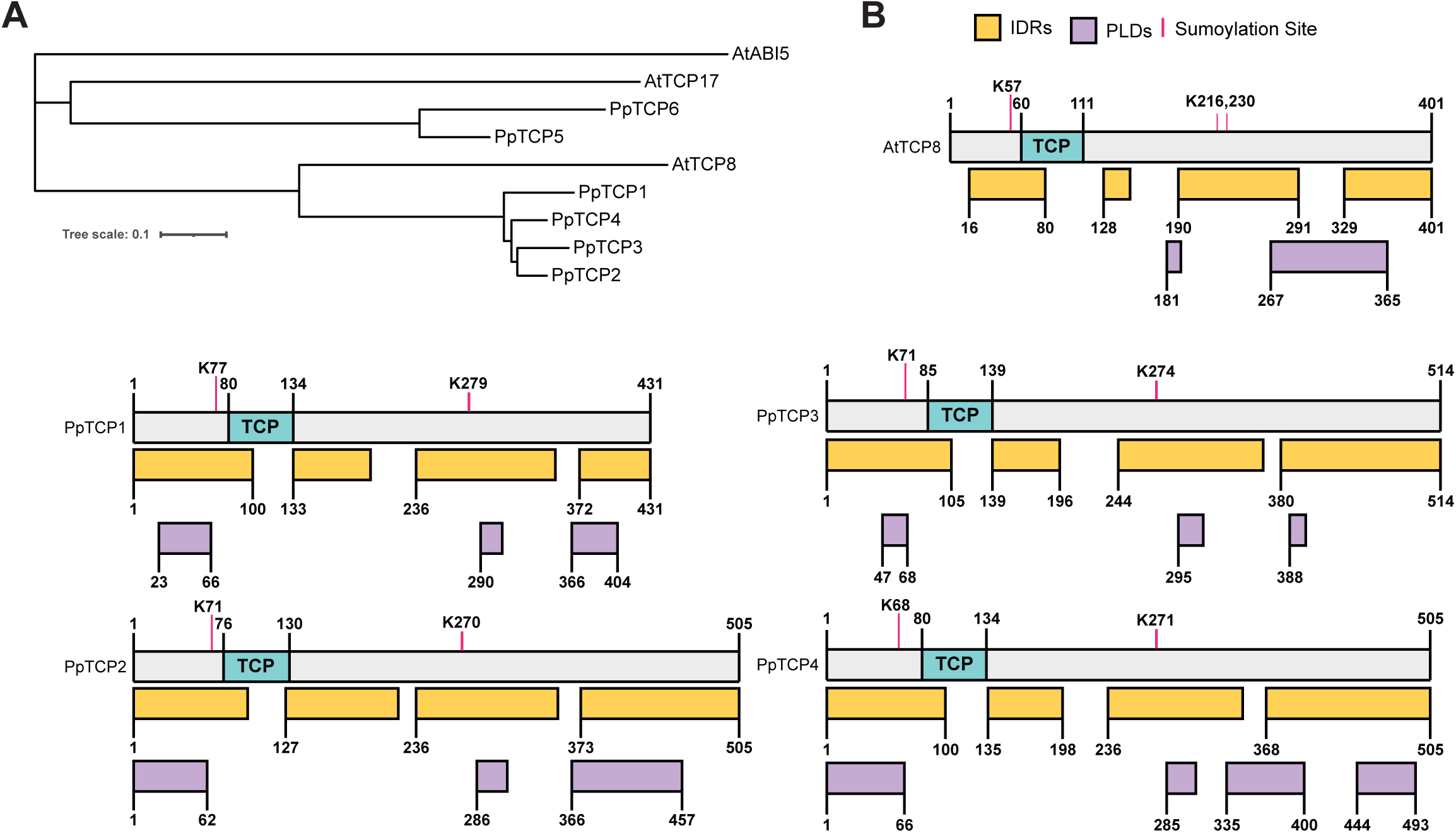
Structural disorder and sumoylation site motifs are conserved between *A. thaliana* and *P. patens* class I TCP-family TFs. **A)** Phylogenetic tree comparing class I PpTCP1-4 and AtTCP8 to class I TCPs and a non-member TF AtABI5. **B)** Similar putative intrinsically disordered regions (IDRs), prion-like domains (PLDs), and sumoylation sites are found in both *A. thaliana* and *P. patens* class I TCPs.

**Figure 4.**
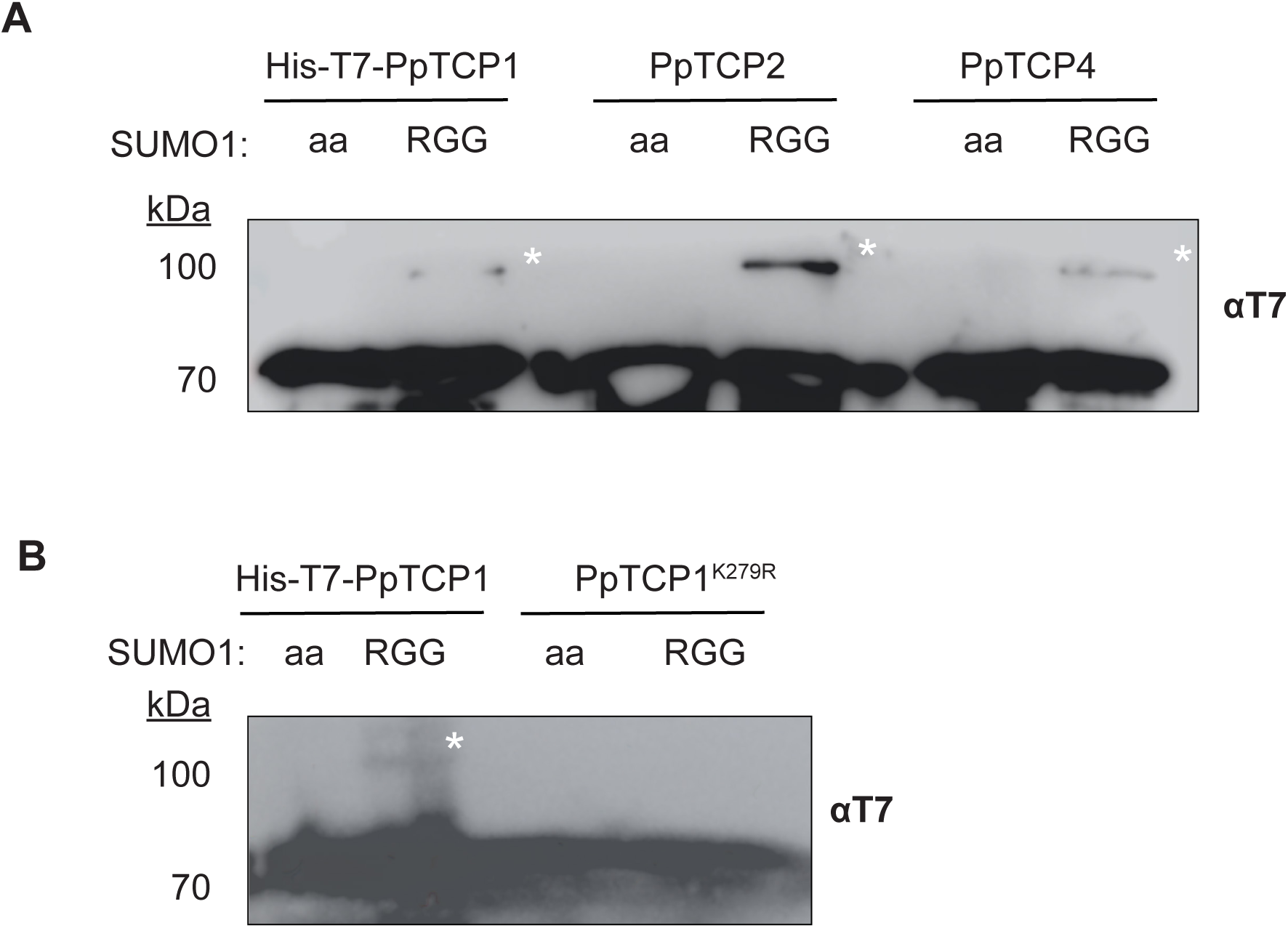
Class I PpTCPs are sumoylated proteins. **A)** Coding regions of class I PpTCPs were cloned into HIS-T7 expression vectors and tested for modification by SUMO1 in the recombinant expression system. All PpTCPs tested exhibited a band shift in the presence of RGG sumo isoforms, indicating sumoylation. **B)** The K279R mutation is sufficient to attenuate the band shift (asterisk) caused by sumoylation of HIS-T7-PpTCP1 by HIS-SUMO1-RGG in the recombinant expression system. Western blots used a T7-directed antibody. Similar results were obtained in three experimental replicates.

### AtTCP8 sumoylation site mutants fail to complement BR-related mutant phenotypes

We have long suspected that the function of TCP8 in the plant response to BR may depend on its sumoylation status. Accordingly, we generated transgenic native promoter-driven *pAtTCP8:TCP8^K57,230R^-HA* Arabidopsis lines in the *tcp8/tcp14/tcp15* mutant background that is compromised in exogenous 24-epiBL responses. Two independent lines were selected for protein levels comparable to the AtTCP8-HA complementation line (Fig S3) and then tested for altered BR-responsive physiology in a standard root growth inhibition assay with 100 nM 24- epiBL. Both independent AtTCP8^K57,230R^-HA lines failed to complement the known long-root and BR-insensitive phenotype of the *tcp8/tcp14/tcp15* mutant, while the functional TCP8-HA complement fully restored BR sensitivity to WT levels (Fig. 5). These results suggest that sumoylation of AtTCP8 positively regulates its activities in promoting BR signaling in Arabidopsis.

**Figure 5.**
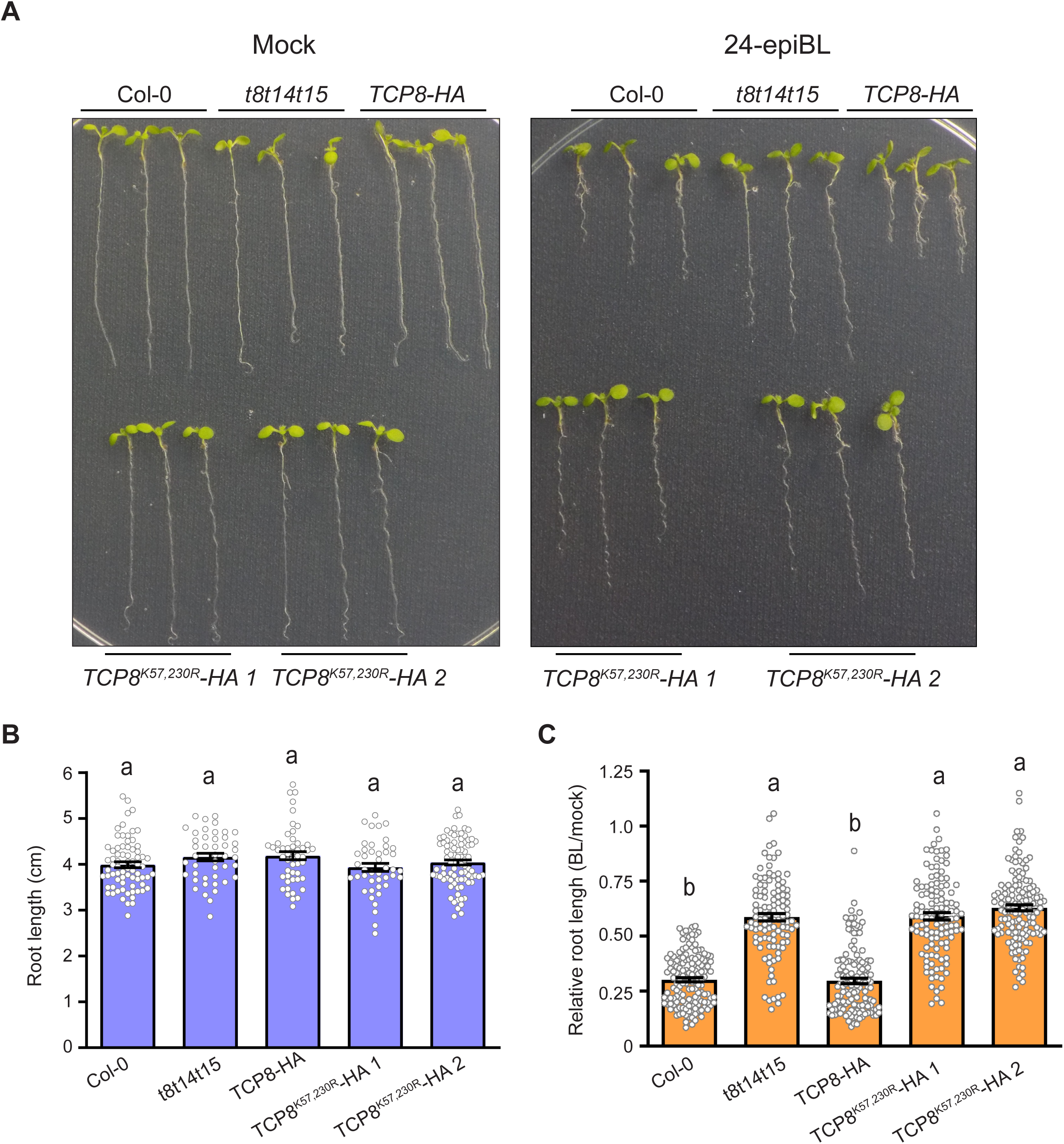
AtTCP8 sumoylation sites regulate BL sensitivity. **A)** A representative plate of seedlings grown vertically on 1/2 MS plates containing DMSO (mock) or 100 nM 24-epiBL for 10 days. **B)** Mock-treated seedling root lengths of Col-0, *tcp8tcp14tcp15*, a triple mutant *pTCP8:TCP8-HA* complementation line, and two different triple mutant *pTCP8:TCP8^K57,230R^-HA* complementation lines were measured using ImageJ. No significant difference in root length was observed between any of the lines tested, as determined by ANOVA with Tukey MCS, p< 0.01, n > 60. Bars are 1 S.E. **C)** Root lengths were determined in 24-epiBL-treated seedlings and normalized to mock. The *TCP8^K57,230R^* complementation lines exhibited BL insensitivity with reduced shortening of roots relative to the WT seedlings, and indistinguishable from the triple mutant. Significance determined by ANOVA with Tukey MCS, p< 0.01, n >108. Bars are 1 SE.

To test this idea, we determined the effects of loss of the K57 and K230 residues on AtTCP8 activity at the BZR2 promoter, a known regulatory target. We transiently expressed *35S:HA- AtTCP8^K57,230R^* with the *pBZR2:GUS* reporter construct in *N. benthamiana* leaf epidermal cells to quantify AtTCP8 activity levels and found that the sumoylation site mutant form of the protein is significantly less active at the promoter region than WT protein, supporting our phenotypic data in Arabidopsis (Fig. 6A). Since AtTCP8 is known to dimerize with AtBZR2 in its regulatory activities, potentially as a feedback mechanism for induction of AtBZR2 expression, we further explored interactions between AtTCP8^K57,230R^ and BZR2 at the protein level using a split luciferase assay in *N. benthamiana*. Interestingly, C-terminally tagged AtTCP8^K57,230R^-HA-cLUC exhibited enhanced interaction with AtBZR2-HA-cLUC relative to WT AtTCP8 (Fig. 6B). These data suggest that AtTCP8 lysines K57 and K230, and by extension the sumoylation of TCP8 at those residues, are likely required for its full activity as part of the *Arabidopsis* BR response.

**Figure 6.**
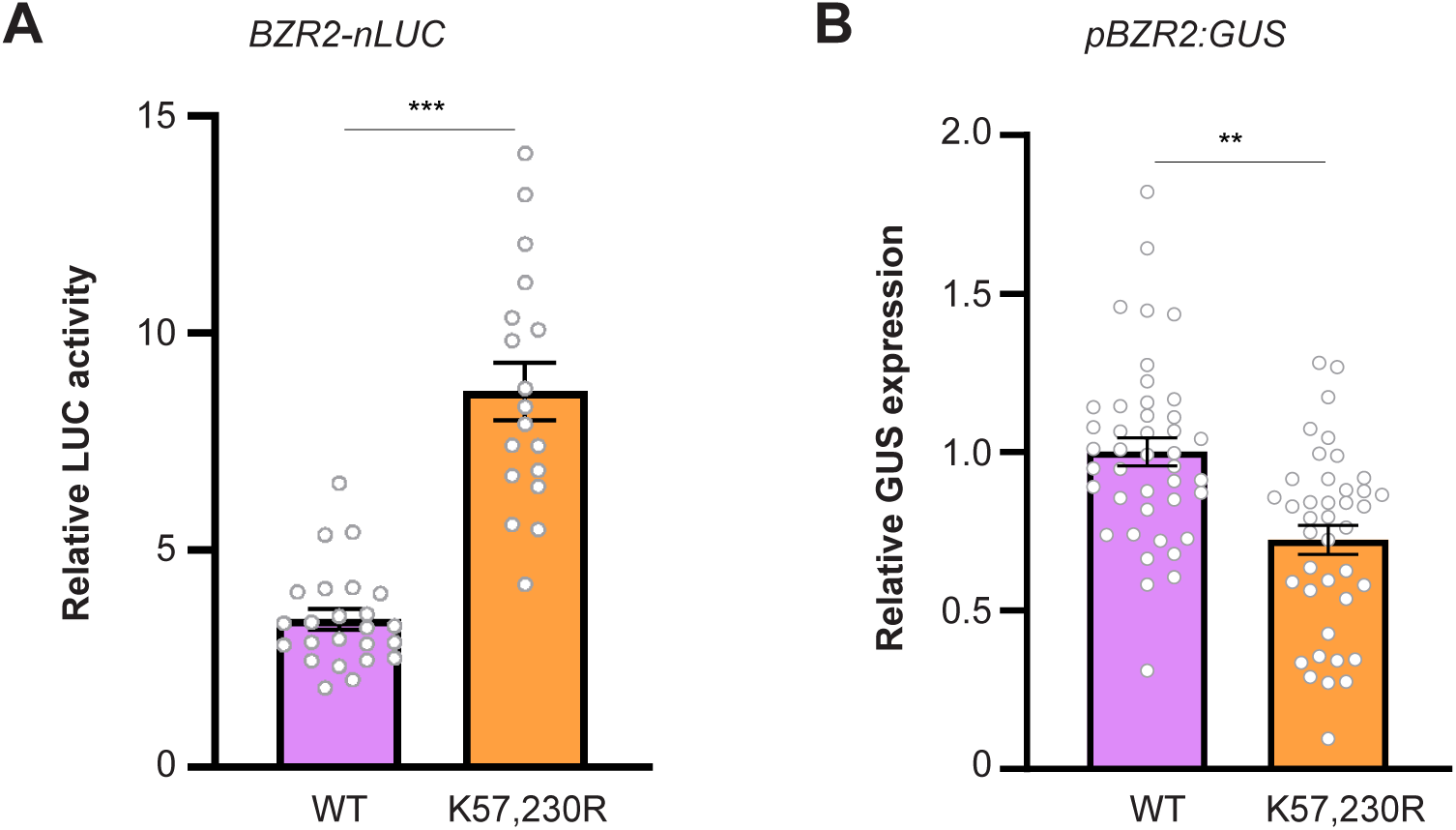
*AtTCP8* activities and dimerization patterns are altered in the *TCP8^K57,230R^* mutant. **A)** Interactions between WT AtTCP8/TCP8^K57,230R^ and known interaction partner AtBZR2 were evaluated by split-luciferase assay in *N. benthamiana* leaf epidermal cells. Total RLUs were normalized to non-interacting control (nLUC-GUS) levels as relative luciferase activity. The K57,230R mutation enhanced interaction with AtBZR2 significantly. n = 20-25 leaf discs. **B)** Transactivation of a BR-responsive reporter gene construct by AtTCP8 is attenuated by K57,230R mutation. Relative GUS expression was determined by normalization to each tested protein’s activation of an EV control. Transcriptional activity of AtTCP8 at the AtBZR2 promoter region was compromised by the K57,230R mutation. n= 40-42 leaf discs. In both experiments, Error bars indicate 1 SE, significance determined by Student’s *t* test at **P <0.01, ***P<0.001

Further, these same residues appear significant to limiting inhibition of AtTCP8 through dimerization with BR-signaling proteins like AtBZR2.

## Discussion

Post-translational modification of plant transcriptional regulators is an effective evolutionary strategy to increase depth of transcriptional output and proteome complexity (Chen *et al*., 2021). A regulator that can adopt multiple conformations under different conditions can assume many different cellular roles. The involvement of AtTCP8 and other disordered class I TCP-family transcription factors in most phytohormone signaling networks as highly connected, pathogen-targeted signaling hubs would suggest a requirement for efficient regulatory mechanisms to appropriately optimize gene expression (Weßling *et al*., 2014; Yang *et al*., 2017; Spears *et al*., 2022). Although post-translational modification of TCP transcription factors has begun to be explored, mechanisms of regulation have yet to be elucidated for most family members and most modifications. Here, we have provided evidence supporting a novel mechanism in which the promotion of BR signaling by AtTCP8 depends on post-translational modifications by SUMO peptides at key residues. Mutation of these residues reduces transcriptional activity of AtTCP8 and inhibits complementation of BR-related phenotypes in *tcp8tcp14tcp15* KO lines, potentially through enhanced dimerization with AtBZR2 as negative feedback to repress BR-related transcript production (Fig. 7). This is a common mechanism in phytohormone signaling that may be conserved from nonvascular plants limited in transcriptional complexity by smaller representation of protein families.

**Figure 7.**
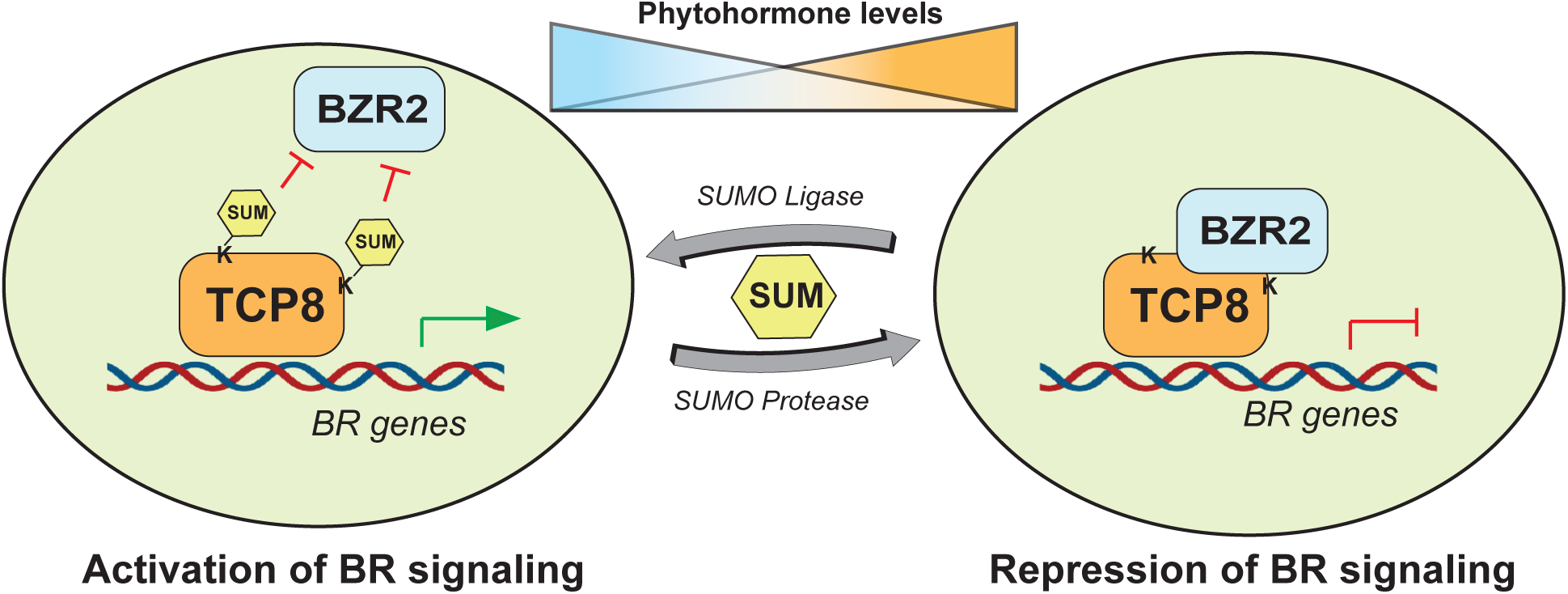
A model describing a potential mechanism for post-translational regulation of AtTCP8 activity through SUMO conjugation. As the levels of phytohormones such as BR fluctuate in the plant cell, TCP8 is reversibly sumoylated or desumoylated. Interaction with latent BZR2 is inhibited when TCP8 is sumoylated at lysines K57 and K230, preventing the inhibition of TCP8 activity at BR-related gene promoters. When desumoylated by SUMO proteases under different conditions, TCP8 is capable of interacting with BZR2, repressing its activity at those promoters.

Our findings that *P. patens* class I TCP proteins are capable of being sumoylated in a recombinant system supports this potential regulatory mechanism in bryophytes, but questions remain. Surely this mechanism is not limited to BR signaling; after all, BR-related genes are only a small subset of the regulatory targets for AtTCP8, and it is yet inconclusive if brassinolide biosynthesis or perception occurs in moss, although different types of BRs have been identified (Yokota *et al*., 2017) and related receptor elements from *P. patens* have been demonstrated to complement *bri1* mutant functionality in *Arabidopsis* (Zheng *et al*., 2019). It would be interesting to see if the sumoylation of AtTCP8 and PpTCPs is also relevant to other phytohormone signaling responses, such as established roles in SA-mediated immune signaling, or suggested involvement in the plant response to auxin in *Arabidopsis*. It is also possible that the previously suggested role of AtTCP8, and now the PpTCPs not as primary regulators of any one phytohormone pathway, but rather as a baseline regulator of phytohormone balance across pathways through dimerization potential with other TFs is an attractive explanation. Sumoylation has already been described in the regulation of immunity-related TFs like AtWRKY33 (Verma *et al*., 2021), and sumoylation machinery mutants have been characterized for compromised immune signaling (Lee *et al*., 2007; van den Burg and Takken, 2010; Ingole *et al*., 2021).

Conjugation of SUMO may represent an antagonistic modification to either phosphorylation or O-GlcNAcylation, sites for which have been identified in the immediate vicinity of characterized sumoylation sites in this study, noticeably clustered around predicted IDRs (Xu *et al*., 2017) (Fig S4). Distinct combinations of post-translational modification could temporarily stabilize the structure of AtTCP8 in response to environmental stimuli, resulting in altered subcellular localization patterns, dimerization habits, and function. In this way, a limited pool of class I TCPs could be capable of governing a wide swath of phytohormone signaling transcriptional output. Further characterization of this family and such mechanisms of regulation will be key to understanding their broad roles in plant growth and development.

## Author Contributions

B.J.S and W.G. planned and designed the research; B.J.S., R.M., S.A.M, N.M.B., N.C.B., and Q.V. performed experiments; B.J.S, R.M., S.A.M., N.A.M, N.C.B, Q.V., and W.G. analyzed and interpreted data; B.J.S., S.A.M., and W.G. wrote the article with editing contributions

