## Supplemental for "Regulation of brassinosteroid signaling through Arabidopsis transcription factor TCP8 by a conserved post-translational mechanism"

**A**

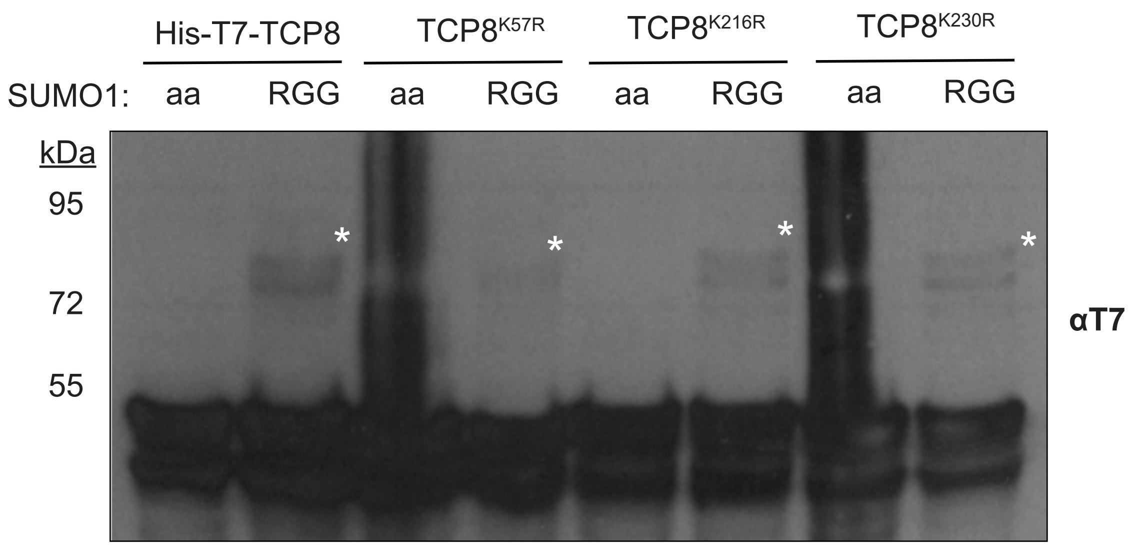


**B**

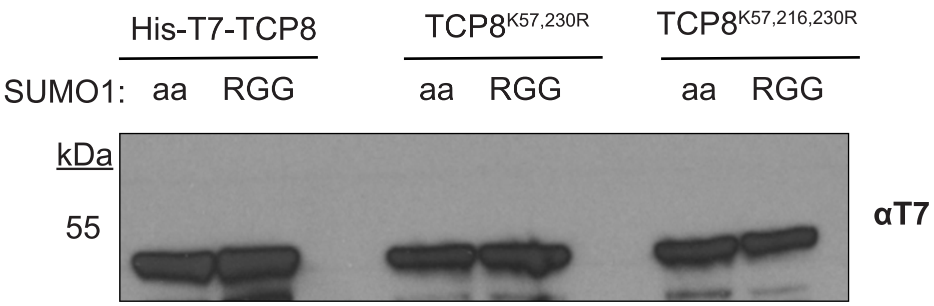


**Figure S1.** In vitro sumoylation assays with AtTCP8$^{KxR}$ mutants. **A)** Sumoylation of AtTCP8 in the recombinant expression system is not attentuated by single K57R, K216R, or K230R mutations. Similar results were observed in two repeat experiments. **B)** Lower exposure image of Figure 2 showing comparable expression levels of HIS-T7-AtTCP8 variants.

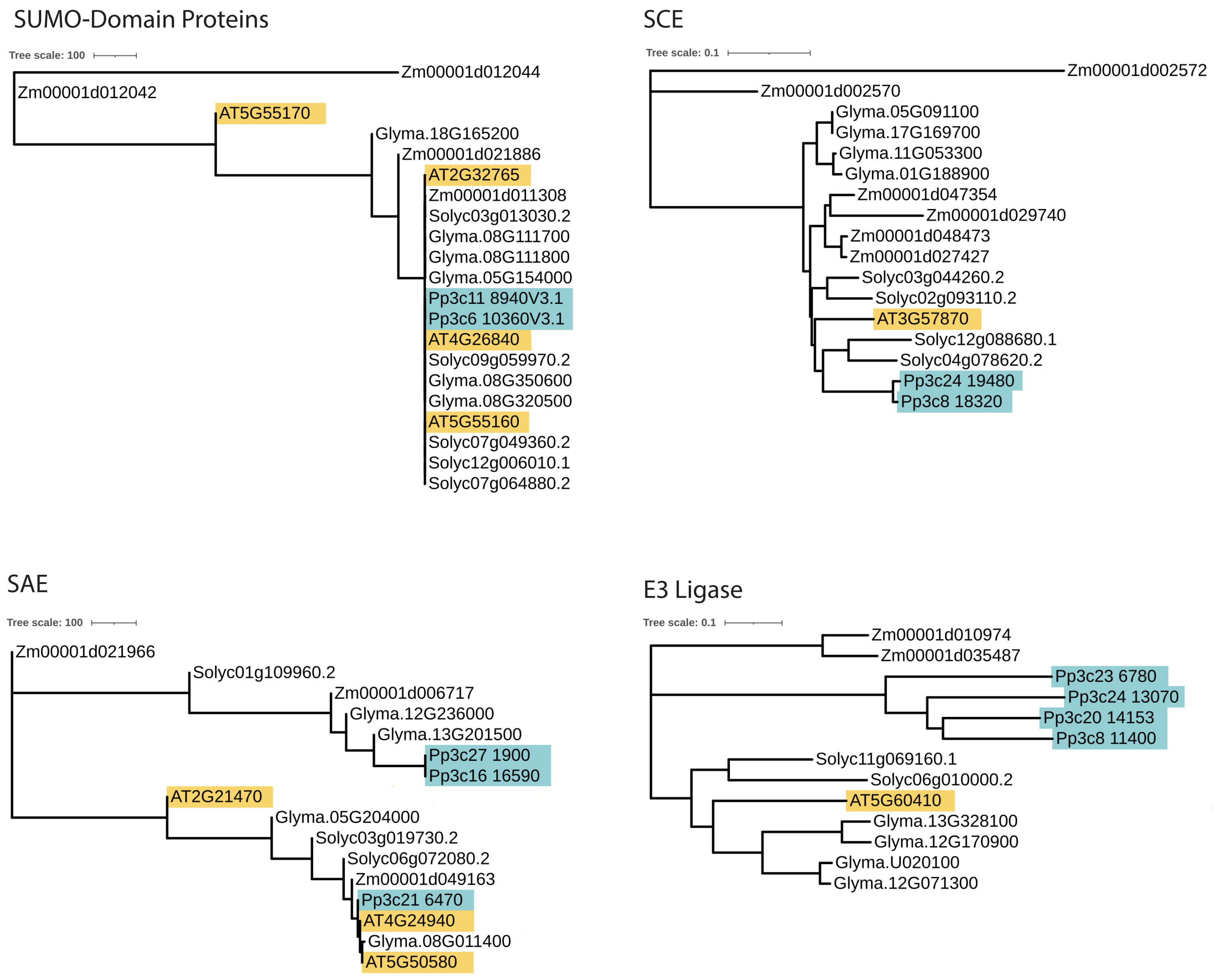


**Figure S2.** Sumoylation machinery is conserved between vascular and non-vascular plant species. *A. thaliana* SUMO Activating Enzyme (SAE)  SUMO protein, SCE, and E3 Ligase sequences are highlighted in orange and *P. patens* sequences are highlighted in blue.

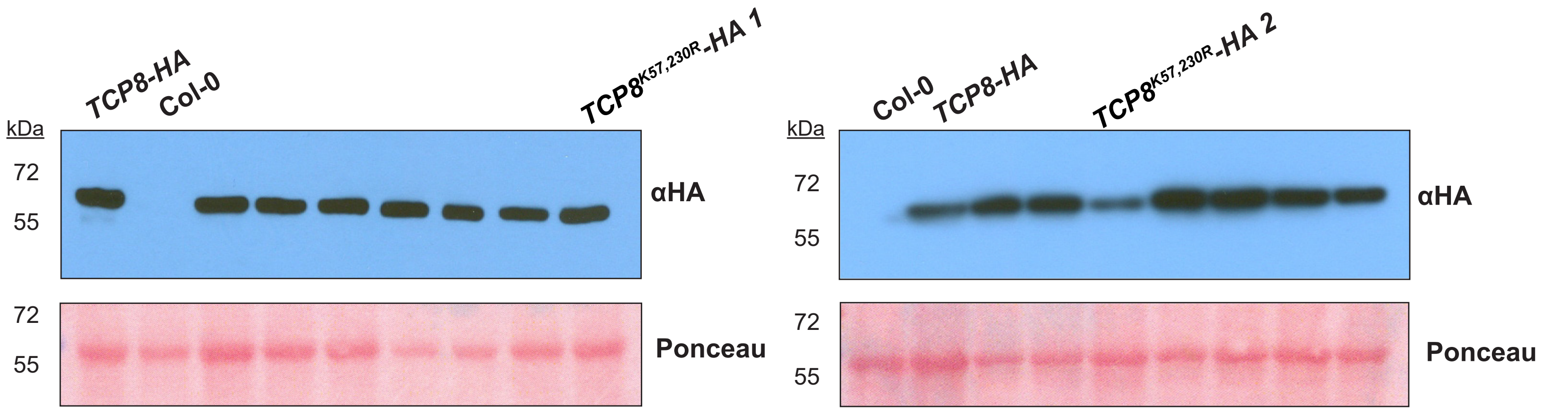


**Figure S3.** *TCP8$^{K57,230R}$-HA* complementation line screen for WT expression levels**.** The *tcp8 tcp14 tcp15* triple mutant was transformed with a *pTCP8:TCP8$^{K57,230R}$-HA* expression contruct and T3 homozygous lines were obtained. For each line tested, four 10-day old *Arabidopsis* seedlings were ground in 150 µl of lysis buffer, 30 µl of which was loaded onto 8% SDS-PAGE gels. Protein was detected with αHA antibody and normalized by ponceau staining of the membrane. The two *TCP8$^{K57,230R}$* lines (1 and 2)in Figure 5 were selected for exhibiting expression levels similar to the WT complentation line.

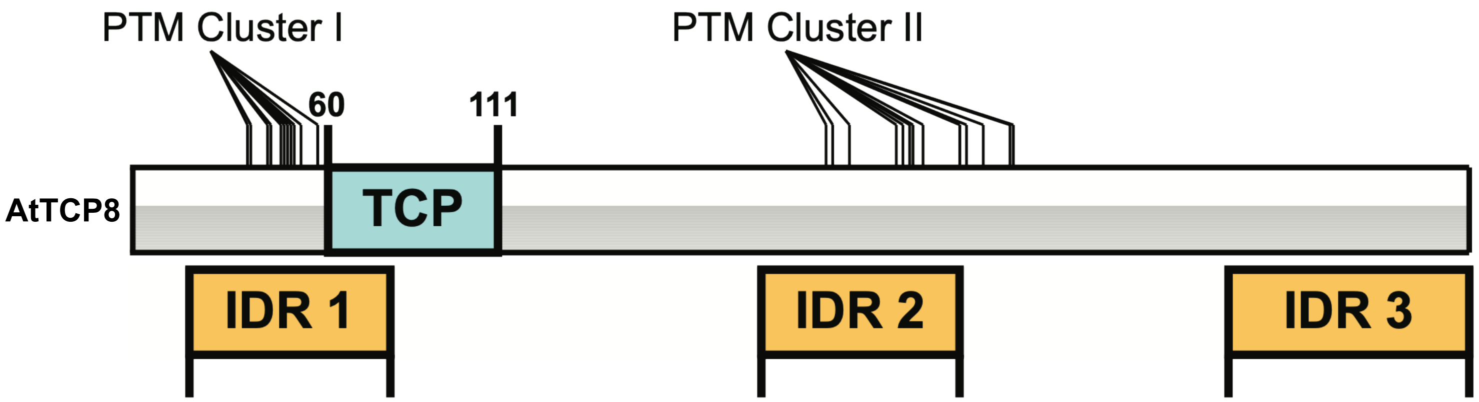


**PTM**
SUMO: K57,216,230
O-GlcNAc: S36,42,48,49,232,235,249,251; T37,43,46,47,50,52, 234,238,256,264,265 (Xu et al, 2017)
Phosphorylation: S36,42,48,49; T37,43,46,47,50; Y206 (Xu et al, 2017)

**Figure S4.** Locations of known post-translational modifications of AtTCP8**.** To date, sites for modification by O-GLcNAc, Phosphate, and SUMO grops have been identified clustered in IDRs near the N-terminal TCP domain and C-terminus of the protein.
